# Arabidopsis Class I Formin FH1 Relocates Between Membrane Compartments During Root Cell Ontogeny And Associates With Plasmodesmata

**DOI:** 10.1101/419853

**Authors:** Denisa Oulehlová, Eva Kollárová, Petra Cifrová, Přemysl Pejchar, Viktor Žárský, Fatima Cvrčková

**Affiliations:** Department of Experimental Plant Biology, Faculty of Science, Charles University, CZ 128 43 Prague 2, Czech Republic; Institute of Experimental Botany of the CAS, CZ 165 02 Prague 6, Czech Republic

**Keywords:** *Arabidopsis thaliana*, At3g25500, cytoskeleton, endomembrane system, plasmodesmata, vacuole

## Abstract

Formins are evolutionarily conserved eukaryotic proteins engaged in actin nucleation and other aspects of cytoskeletal organization. Angiosperms have two formin clades with multiple paralogs; typical plant Class I formins are integral membrane proteins that can anchor cytoskeletal structures to membranes. For the main Arabidopsis housekeeping Class I formin, FH1 (At3g25500), plasmalemma localization was documented in heterologous expression and overexpression studies. We previously showed that loss of FH1 function increases cotyledon epidermal pavement cell shape complexity via modification of actin and microtubule organization and dynamics. Here we employ transgenic Arabidopsis expressing green fluorescent protein-tagged FH1 (FH1-GFP) from its native promoter to investigate *in vivo* behaviour of this formin using advanced microscopy techniques. The fusion protein is functional, since its expression complements the *fh1* loss-of-function mutant phenotype. Accidental overexpression of FH1-GFP results in a decrease in trichome branch number, while *fh1* mutation has the opposite effect, indicating a general role of this formin in controlling cell shape complexity. Consistent with previous reports, FH1-GFP associates with membranes. However, the protein exhibits surprising actin- and secretory pathway-dependent dynamic localization and relocates between cellular endomembranes and the plasmalemma during cell division and differentiation in root tissues, with transient tonoplast localization at the transition/elongation zones border. FH1-GFP also accumulates in actin-rich regions of cortical cytoplasm and associates with plasmodesmata in both the cotyledon epidermis and root tissues. Together with previous reports from metazoan systems, this suggests that formins might have an ancestral role at cell-cell junctions.

## Introduction

Shaping of plant cells, tissues and organs relies upon precise temporal and spatial control of cell division and cell expansion. These processes depend on orchestrated dynamics of the two cytoskeletons present in plant cells – the microfilaments and microtubules – and the endomembrane system that is ultimately responsible for targeted exocytosis of materials required for cell wall construction. Despite major recent progress (see e.g. Lehman et al. 2017), our understanding of the mechanisms linking the two cytoskeletal systems to each other and to membranes remains far from complete.

Formins, a family of cytoskeletal organizers commonly engaged in modulating both actin and microtubule dynamics and often associating with membranes, are obvious candidates for co-ordinating the cytoskeletons and membranes in the plant cell cortex. All formins share the evolutionarily conserved FH2 domain whose dimer can nucleate and cap actin filaments and which is usually accompanied by diverse additional domains (see Grunt et al. 2008, Breitsprecher and Goode 2013). Many formins directly interact with microtubules both in opisthokonts and in plants (reviewed in Chesarone et al. 2010, Wang et al. 2012). Also membrane association is a widespread feature of formins, although its mechanisms vary. Opisthokont formins are often regulated by small GTPases (which themselves are peripheral membrane proteins), and some non-angiosperm formins contain a possible small GTPase-binding domain. However, small GTPase binding is not predicted for angiosperms, whose formins can be classified into two clades based on FH2 domain phylogeny. While typical Class I formins are integral membrane proteins, Class II formins commonly harbour a phospholipid-binding domain related to the PTEN oncogene, and alternative means of membrane attachment mediated by interacting proteins might be present in formins lacking the typical membrane insertion or membrane-binding motifs (see Cvrčková 2013).

In all angiosperms studied so far, both formin clades are represented by many paralogs; *Arabidopsis thaliana* has 11 Class I and 10 Class II formin-encoding genes (Grunt et al. 2008). The biological role of this diversity, possibly further increased by alternative splicing and/or formin heterodimerization, remains unclear. Especially in case of Class I formins, which typically exhibit a N-terminal Pro-rich extracytoplasmic domain that can enable formin-mediated anchoring of the cortical cytoskeleton to the cell wall (Cvrčková 2000, Martiniere et al. 2011, van Gisbergen and Bezanilla 2013, Borassi et al. 2016), it is tempting to speculate about evolutionary optimization of individual paralogs for functions specific to distinct temporal or spatial domains within the plant body and individual cells. Indeed, restricted lateral mobility in the plasmalemma has been documented for Arabidopsis Class I formin FH1 (Martiniere et al. 2012). Here we shall further focus only on Class I formins and on the *Arabidopsis thaliana* model system, unless stated otherwise.

Distinct Class I formins may exhibit paralog-specific gene expression patterns and/or intracellular localization. The former has been documented in genome-wide expression analyses (Hruz et al. 2008) and by several studies focusing on single genes. Expression of FH6 is induced upon nematode infection (Favery et al. 2004), and FH5 is expressed at the posterior endosperm pole in the developing seed (Ingouff et al. 2005). However, there is only fragmentary evidence for targeting of distinct formin paralogs to specific locations of the plasmalemma or to other membrane compartments. Two closely related formins FH4 and FH8 were found to accumulate at cell to cell borders perpendicular to root axis based on both immunolocalization of native protein and localization of a green fluorescent protein (GFP)-tagged non-functional deletion derivative that retained the N-terminal membrane-targeting signal (Deeks et al. 2005). Recently, both GFP-tagged, biologically active FH4 and its C-terminally truncated derivative lacking the actin-binding domains was found to rapidly relocate to sites of callose deposition in plant cells attacked by fungal or oomycete pathogens (Sassman et al. 2018). Other deletion derivatives of FH4 incapable of membrane insertion associated i*n vivo* with microtubules and the endomembrane system (Deeks et al. 2010); the membrane localization may have been mediated by protein interactions, including possibly dimerization with intact endogenous protein. FH6, as detected by immunolocalization, decorates the plasmalemma of nematode-induced giant cells uniformly, and overexpresed GFP-tagged FH6 was uniformly distributed across the plasmalemma of protoplasts (Favery et al. 2004). Overexpressed, GFP-tagged FH5 localizes to the nascent cell plate during cytokinesis (Ingouff et al. 2005) and is enriched in the apical cytoplasm (or some fine structures therein) of growing pollen tubes (Cheung et al. 2010), while its deletion derivative incapable of membrane insertion accumulates in the endoplasmic reticulum and nuclear membrane (Cvrčková et al. 2014). With the sole exception of FH2, whose GFP-tagged derivative, which complemented a mutant defect in the control of intercellular trafficking, was very recently reported to localize to the plasmodesmata when expressed from the endogenous promoter (Diao et al. 2018), current knowledge regarding Class I formin localization comes from immunodetection (with the possibility of artefacts), from heterologous expression or overexpression experiments involving tagged proteins of unclear functional status, or from studies employing non-functional deletion derivatives. This has been so far also the case of FH1, the Class I formin most ubiquitously expressed in the Arabidopsis sporophyte but barely detectable in the gametophyte (Hruz et al. 2008), which is the topic of the present study.

FH1 (At3g25500) is a typical Class I formin whose ability to stimulate actin polymerization and bundling has been documented *in vitro* (Michelot et al. 2005). Heterologous overexpression in tobacco pollen tubes caused formation of massive actin bundles (Cheung and Wu 2004). While this may suggest that impaired FH1 function should result in reduced actin bundling, microfilaments appear to be actually more bundled and less dynamic in the rhizodermis and cotyledon epidermis of loss-of-function *fh1* mutants, while microtubule mobility is increased (Rosero et al. 2013, Rosero et al. 2016, Cvrčková and Oulehlová 2017). This suggests a more complex relationship, and possibly a fine balance, between FH1 and other formin paralogs. Membrane localization, predicted from the FH1 sequence, was confirmed by cell fractionation in the first published study of plant formins (Banno and Chua 2000). When GFP-tagged and heterologously expressed in tobacco pollen tubes, FH1 uniformly decorated the apical dome plasmalemma, plasmalemma invaginations caused by its overexpression, and possibly some endomembrane compartments (Cheung and Wu 2004). When heterologously expressed in mature *Nicotiana benthamiana* leaves, GFP-tagged FH1 is located at the plasmalemma but excluded from the regions adjacent to underlying cortical microtubules, while a deletion derivative lacking most of the cytoplasmic domains labels the plasmalemma uniformly (Martiniere et al. 2011). However, a longer C-terminally truncated version of the protein was found at the plasmodesmata together with FH2 when expressed in Arabidopsis using the strong constitutive 35S promoter (Diao et al. 2018). Thus, while there is a general consensus on the predominant localization of FH1 at the plasma membrane, some uncertainties due to possible effects of heterologous and/or ectopic expression, overexpression, and possible altered functionality of tagged fusion proteins persist.

We previously found that *fh1* loss-of-function mutants display a subtle, yet reliable and easily detectable phenotype – increased complexity of cotyledon pavement cell shape (Rosero et al. 2016). Here we use complementation of this mutant phenotype to prove functionality of GFP-tagged FH1 construct expressed from the FH1 promoter to achieve close to native protein levels, and characterize localization of this construct in detail, revealing a surprisingly complex pattern of formin relocation between a variety of membrane compartments and domains.

## Results

### GFP-tagged FH1 is biologically active

To examine FH1 localization under conditions as close to native as possible, we prepared stable transgenic *A. thaliana* lines carrying a construct with C-terminally GFP-tagged FH1 under the control of the FH1 promoter (further referred to as *pFH1:FH1-GFP*). When introduced into *fh1-1* mutant plants previously shown to exhibit increased lobing (i.e. decreased circularity) and enlarged size of cotyledon epidermal pavement cells (Rosero et al. 2016), the construct fully complemented both mutant defects, thus confirming biological activity of *pFH1:FH1-GFP* (Fig. 1 A,B). The resulting complemented transgenic plants will be further referred to as *fh1-1c*. Transformed lines were also prepared in a genetic background carrying an independent loss-of-function allele of FH1, *fh1-4,* as well as in a background that was wild type (WT) with respect to *FH1* but carried a *rdr6* mutation to prevent transgene silencing (Sasse et al. 2015). The latter transformants were subsequently used to construct marker or mutant plant lines carrying *pFH1:FH1-GFP* by crossing.

**Figure 1.**
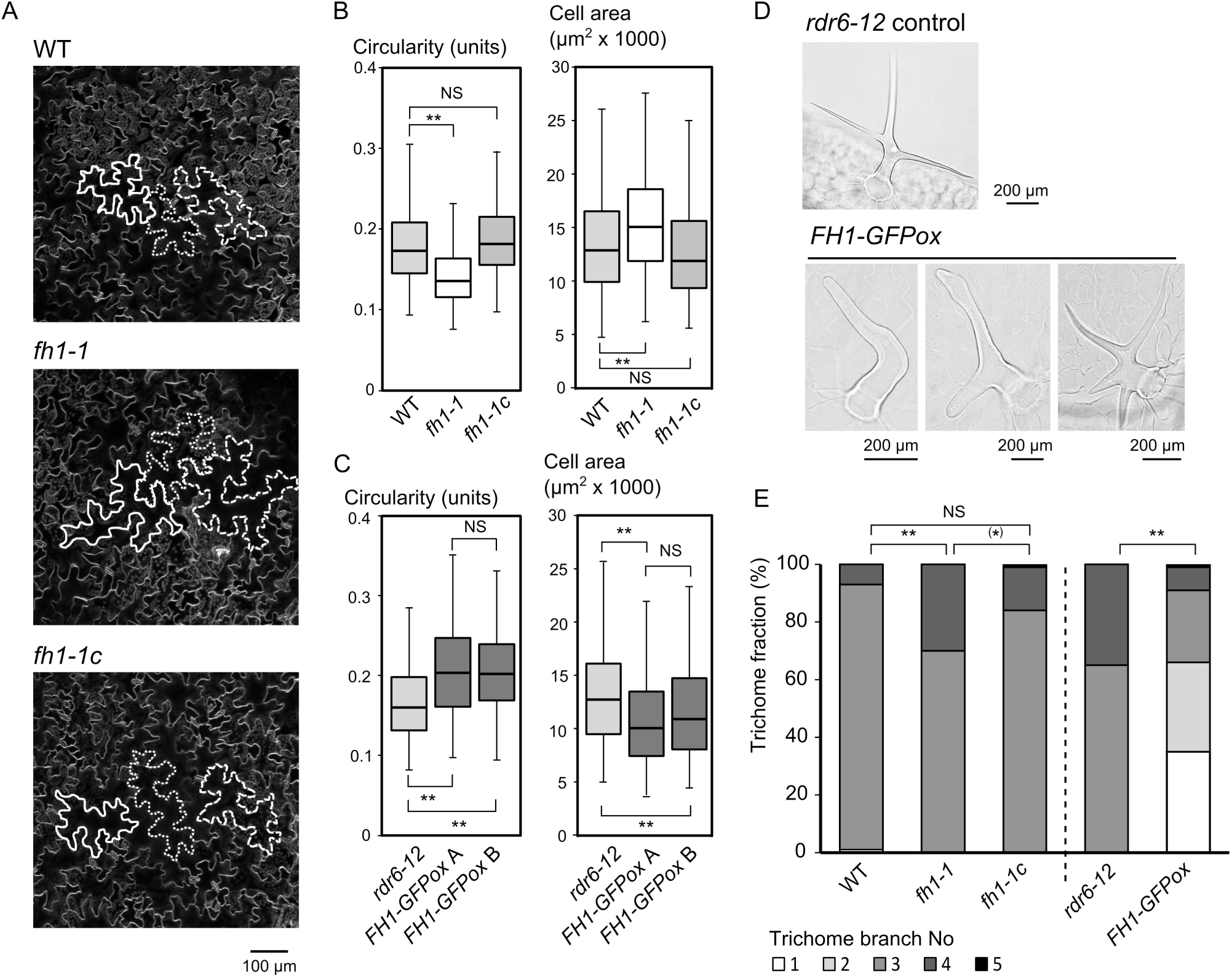
Biological activity of *pFH1:FH1-GFP* documented by complementation of *fh1* and by overexpression.(A) Pavement cells on the adaxial side of cotyledons from 14 days old WT, *fh1-1* mutant and *fh1-1 pFH1:FH1-GFP* (*fh1-1c*). seedlings. (B) Shape parameters of cotyledon pavement cells from 14 days old WT, *fh1-1* and *fh1-1c* seedlings. (C) Shape parameters of cotyledon pavement cells of 14 days old parental *rdr6-12* line and two different overexpression lines originating from the same mother plant (*FH1-GFPox*). In (B) and (C), at least 250 cells per line were analyzed, double asterisks represent significant between-genotype differences (ANOVA, p < 0.01), NS – non-significant. (D) Typical trichomes on fully expanded first true leaves of 3 weeks old *rdr6-12* and *FH1-GFPox* plants. (E) Distribution of trichome branch numbers from first true leaves for the genotypes indicated. At least 120 trichomes were evaluated for each genotype. Asterisks denote statistical significance of between-genotype differences (pairwise Chi square test with Yates correction for zero-count categories and Bonferroni correction for multiplicity, ** – p < 0.01, (*) – marginal, i.e. p < 0.1, NS – not significant, i.e. p > 0.1).

While most *pFH1:FH1-GFP* transformants in the *rdr6* background were phenotypically indistinguishable from non-transformed plants and exhibited low fluorescence levels, we obtained one transformed *rdr6* line with extremely strong fluorescence indicating incidental transgene overexpression, further referred to as the *FH1-GFPox* line (Supplementary data Figure S1). These plants exhibited decreased cotyledon pavement cell lobing and size, i.e. a phenotype opposite to that of *fh1* mutants (Fig. 1 C), consistent with biological activity of the FH1-GFP fusion protein.

### FH1 contributes to the control of trichome shape

While observing the *FH1-GFPox* seedlings, we noticed that their first leaves often carry aberrantly shaped trichomes (Fig. 1 D). Although we cannot entirely exclude the possibility that this might be a consequence of a transgene-induced insertional mutation outside the *FH1* locus, presence of this phenotype completely correlated with that of the transgene in the progeny of a heterozygous primary transformant (i.e. the trichome deformity phenotype was dominant). This observation prompted us to examine trichomes of WT, *fh1* mutants and *FH1-GFP* transgenic lines in more detail. While *fh1* mutants had usually a larger fraction of trichomes with more than three branches than WT plants, this effect was compensated in *fh1-1c* transformants. In contrast, *FH1-GFPox* plants had fewer trichome branches (Fig. 1 E).

Thus, besides its known role in pavement cell shaping, FH1 appears to affect the layout of trichome branches, possibly in a manner analogous to its participation in lobe initiation in pavement cells.

### FH1 localization during early epidermal cell morphogenesis

Since pavement cell lobe initiation takes place during early stages of differentiation (Armour et al. 2015), we examined FH1 localization in young, not yet fully expanded cotyledons and leaves. FH1-GFP uniformly decorates newly formed crosswalls, but later the signal becomes restricted to at least two populations of particles – immobile ones located at or near the plasmalemma, and mobile ones within cortical cytoplasm (Fig. 2 A, Supplementary data Video S2 showing particles in the cortical cytoplasm of epidermal pavement cells of young true leaf of an 8 days old *rdr6 pFH1:FH1-GFP* plant, acquired under the same conditions as the images shown in Fig. 2 A; scale bar = 10 µm).

**Figure 2.**
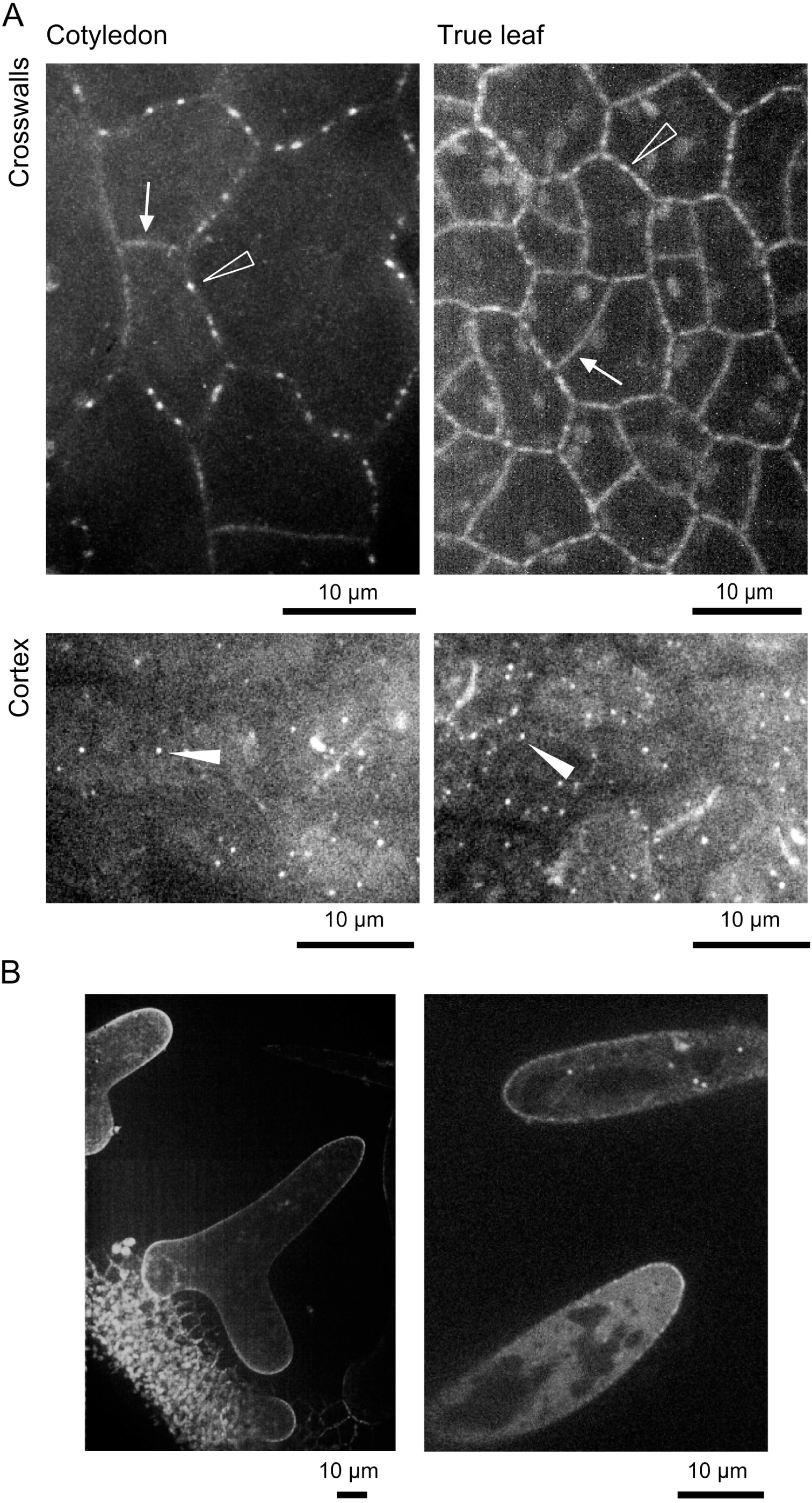
FH1-GFP localization in cotyledon and leaf epidermis. (A) Localization in pavement cells of immature cotyledons (3 days old *fh1-4 pFH1:FH1-GFP* seedlings) and leaves (8 days old, *rdr6 pFH1:FH1-GFP OX* seedlings); single confocal sections focused at the level of anticlinal walls (crosswalls) or periclinal cortical cytoplasm (cortex) are shown. Arrows denote uniformly labelled crosswalls, open arrowheads immobile plasmalemma-associated particles, filled arrowheads cortical particles, often mobile. (B) Z-projections of FH1-GFP distribution in developing trichomes; left – trichome bulges and very young trichomes of a 5 days old *fh1-4 pFH1:FH1-GFP* plant, right – branches of older yet still expanding trichomes of an 8 days old *rdr6 pFH1:FH1-GFP OX* seedling.

During trichome development, starting from the early bulge stage, FH1-GFP localizes predominantly at the tips of growing trichome branches, but some mobile particles were also observed, together with weak cytoplasmic signal that becomes more apparent as a trichome grows (Fig. 2 B). No signal was detected in mature trichomes, consistent with FH1 acting mainly in early trichome development.

### FH1 is targeted to plasmodesmata and co-localizes with actin, but not microtubules, in differentiated pavement cells

In fully differentiated lobed pavement cells the mobile cytoplasmic particles are absent and FH1-GFP localization is restricted to the plasmalemma and immobile dots therein. Plasmalemma localization can be documented by complete co-localization of the FH1-GFP signal with the membrane-staining FM4-64 dye, which persists even after plasmolysis (Fig. 3). Interestingly, the local fluorescence maxima in the plasmalemma weaken upon plasmolysis as the membrane becomes stretched, except in regions where the plasmalemma failed to detach from the cell wall.

**Figure 3.**
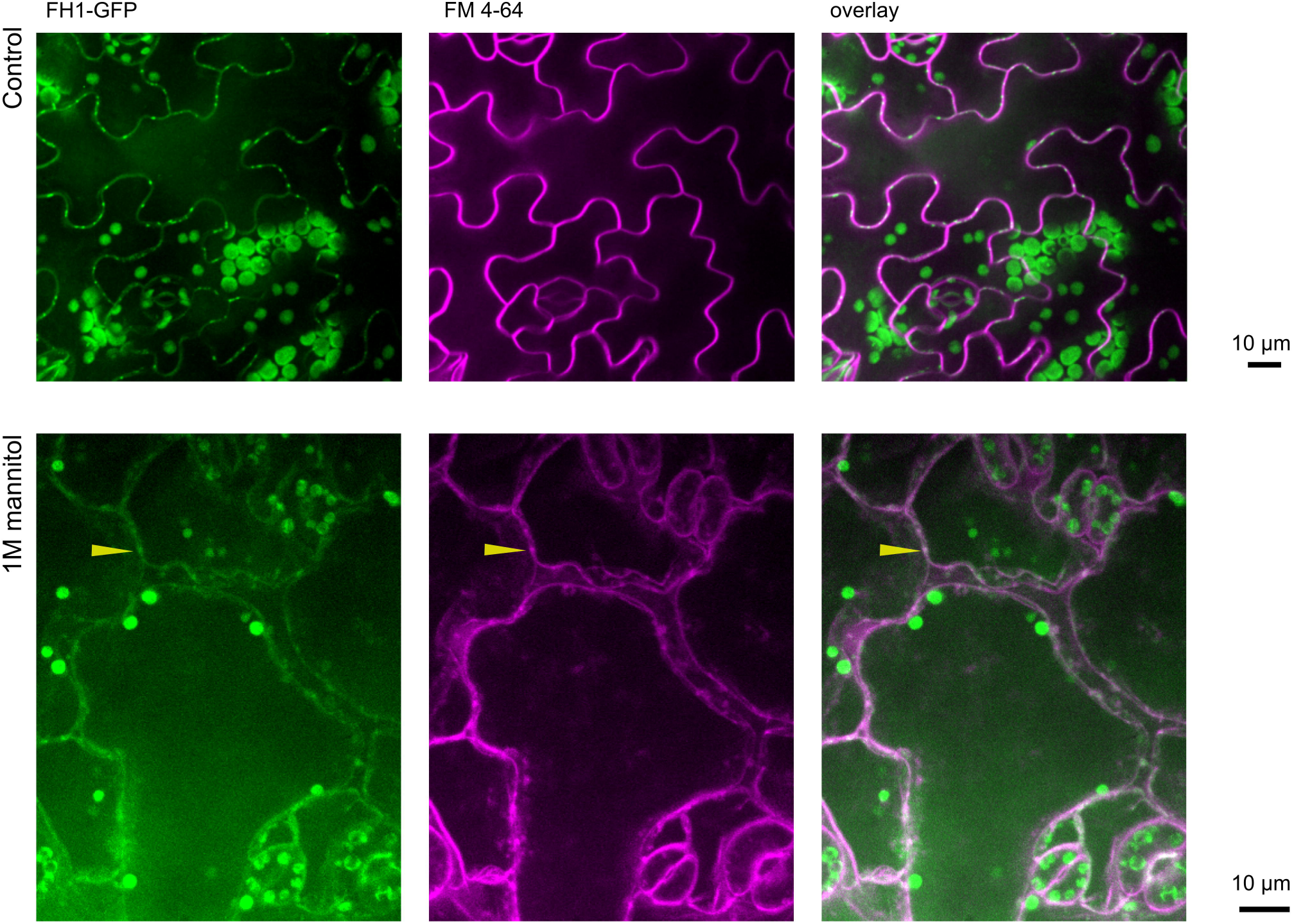
FH1-GFP is targeted to the plasmalemma in pavement cells. Top: Z-projections of FH1-GFP and plasmalemma (FM4-64 staining) in differentiated cotyledon pavement cells of 6 days old *fh1-4 pFH1:FH1-GFP* seedlings. Bottom: FH1-GFP localization after plasmolysis induced by 1M mannitol. Yellow arrowhead points to a preserved FH1-GFP spot at a place where the plasmalemma remained attached during plasmolysis. Some chloroplast fluorescence leaking into the GFP channel is visible.

The punctate plasmalemma localization suggested that FH1-GFP might be associated with plasmodesmata. We confirmed that this is the case, since the formin co-localizes with the receptor-like kinase encoded by the At5g24010 locus which is targeted to plasmodesmata (Fernandez-Calvino et al. 2011) but also labels the plasma membrane in young tissues (Fig. 4, Supplementary data Figure S3 A).

**Figure 4.**
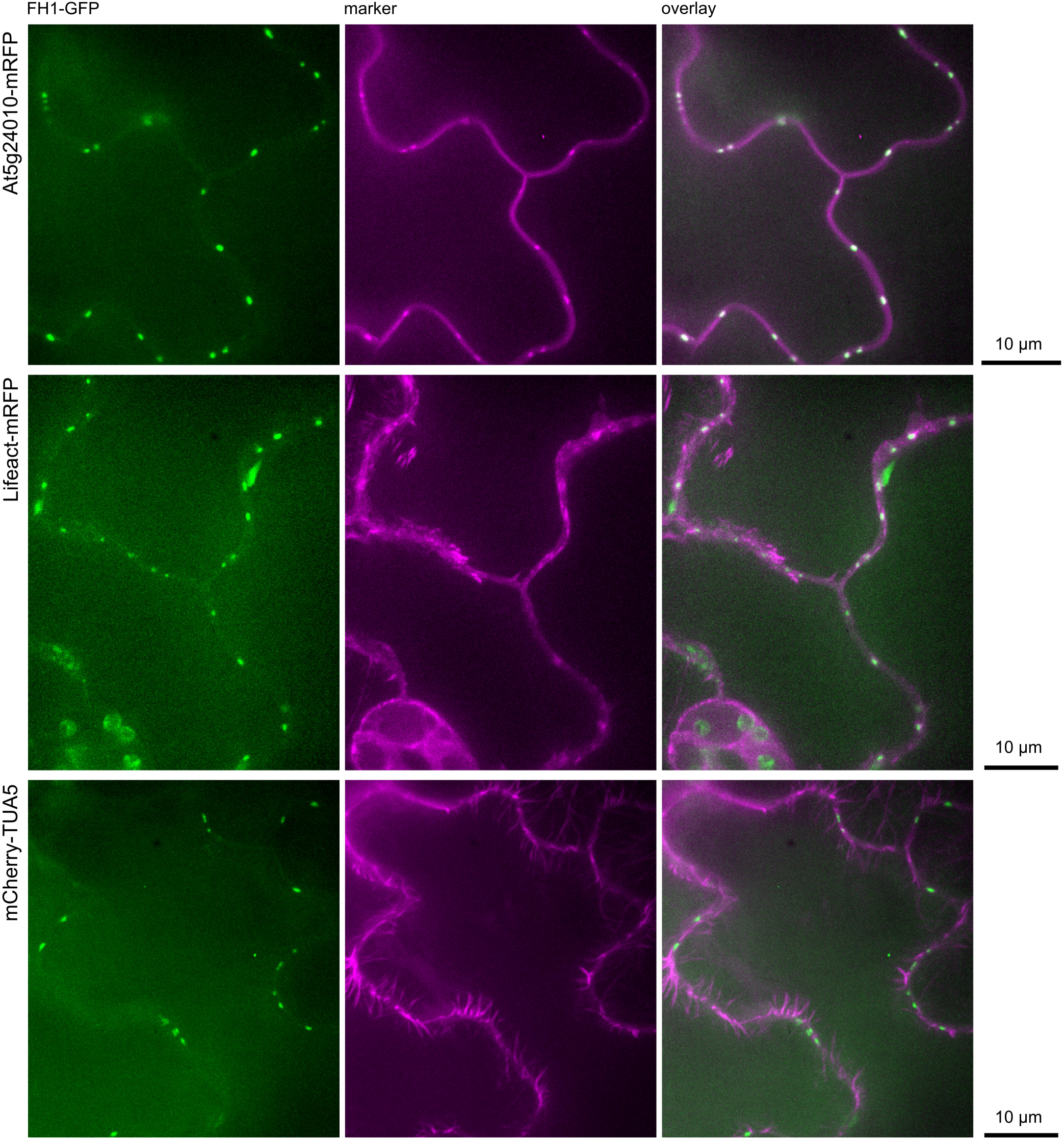
FH1-GFP associates with plasmodesmata and actin but not microtubules. Z-projections of cotyledon epidermis of 6 days old *pFH1:FH1-GFP* seedlings from the progeny of a cross between plants carrying *pFH1:FH1-GFP* in *fh1-4* or *rdr6-12* background and red marker lines. Co-localization of FH1-GFP with RFP-tagged plasmodesmata (At5g24010,) actin (35S:Lifeact-mRFP)and microtubule (35S:mCherry-TUA5) markers is shown.

We also crossed *pFH1:FH1-GFP* transgenic lines to lines carrying cytoskeletal marker proteins. The formin associated with actin-rich cortical domains but not with microtubules, appearing, on the contrary, to be excluded from microtubule-rich cortical regions (Fig. 4, Supplementary Data Figure S3 A), is in agreement with previous observations (Martiniere et al. 2011).

### Complex patterns of FH1 relocation during root ontogeny

In cotyledon pavement cells FH1 relocates between distinct membrane structures (plasmalemma, particles that may correspond to endomembrane compartments, and plasmodesmata) during ontogeny. To explore this phenomenon in more detail, we focused on roots, where cells at various differentiation stages can be observed simultaneously in distinct developmental zones. Consistent with the previously documented peak of FH1 expression in immature root tissues (Brady et al. 2007), fluorescent signal was observed throughout the tip of the main root, but its localization differed among developmental zones.

In the apical meristem, FH1-GFP accumulates at the plasmalemma and in intracellular particles or compartments. The plasmalemma signal appears to be unevenly distributed with enrichment at cell to cell boundaries perpendicular to the root axis (Fig. 5 A,B); the FH1-GFP signal is discernible already at the early cytokinesis stage prior to phragmoplast expansion, associated with the nascent cell plate (Supplementary data Figure S3 B).

**Figure 5.**
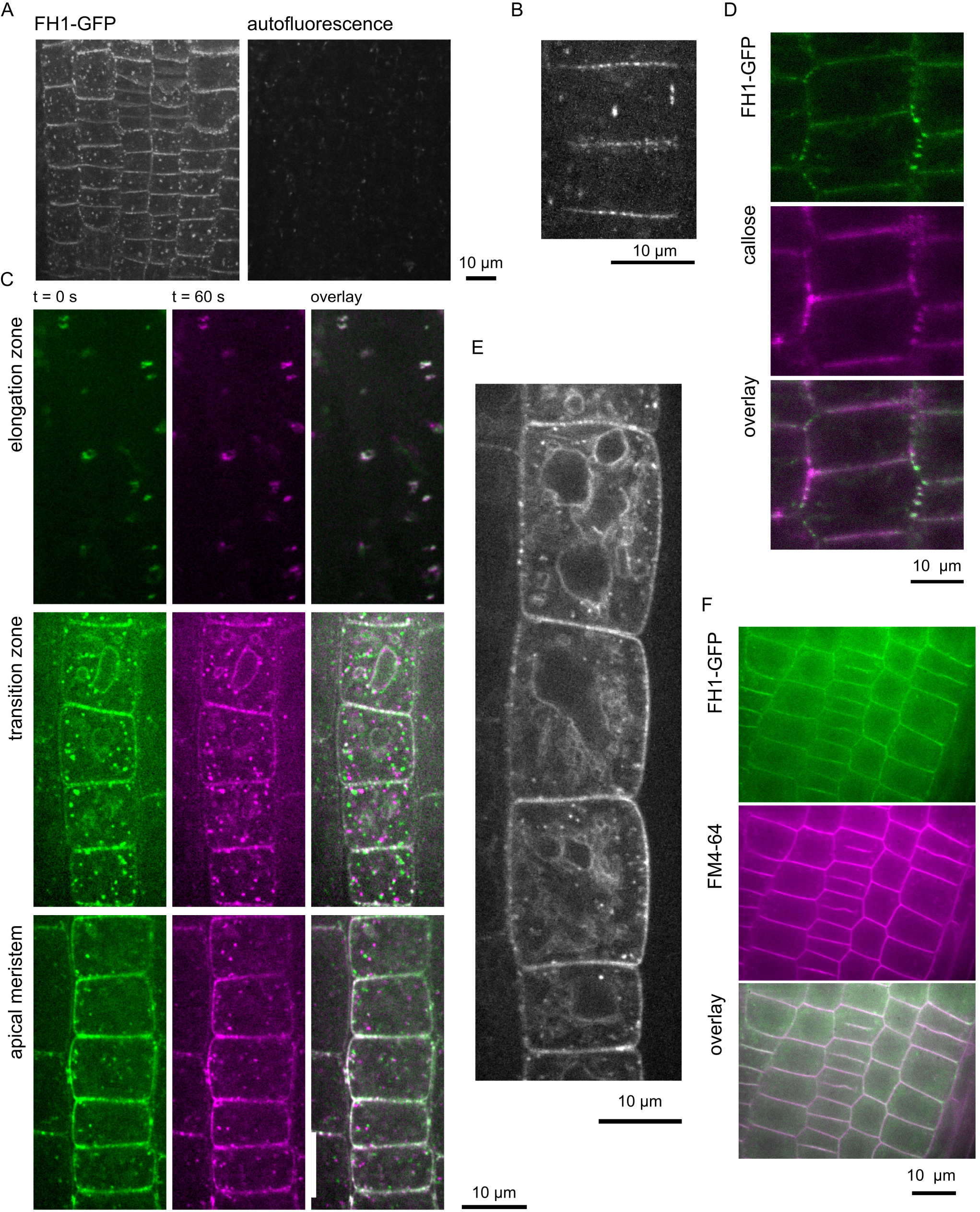
FH1-GFP localizes to various membrane compartments in the root tips of *fh1-1 pFH1:FH1-GFP, fh1-4 pFH1:FH1-GFP* and *rdr6-12 pFH1:FH1-GFP* seedlings. (A) SDCM image of FH1-GFP signal in the apical meristem with autofluorescence control. (B) Higher magnification of the apical meristem, signal accumulates in anticlinal membranes. (C) Single planes from different zones of a root tip taken 60 s apart and their overlays. While the FH1-GFP signal at the plasma membrane remains static (white in the overlays), particles inside cytoplasm are mobile. (D) Co-localization of FH1-GFP and callose in the root meristem (single confocal plane). (E) Z-projection of transition zone showing tonoplast labelling. (F) Apical meristem region stained with FM4-64 (note localization of FH1-GFP to nascent cell plates). All images are from 6 days old seedlings except (D), which is from 4 days old plants.

Similar to the aboveground organs, also in the root apical meristem and in the transition zone cortex FH1-GFP-positive particles can be divided into two populations even within one cell – mobile dots in the cortical cytoplasm and static ones at the plasma membrane, which by analogy with the above ground organs may be associated with plasmodesmata. (Fig. 5 C, Supplementary data Video S4 showing particles in the root apical meristem of a *fh1-4 pFH1:FH1-GFP* plant, acquired under the same conditions as the images shown in Fig. 5 A; scale bar = 10 µm.). Since our plasmodesmal marker At5g24010 uniformly labels the plasma membrane in roots, we employed aniline blue staining of callose to examine co-localization with plasmodesmata in the root tissues. Indeed, the static FH1-GFP foci co-localized with sites of callose deposition, consistent with its plasmodesmatal localization (Fig. 5 D).

Surprisingly, in a subset of transition zone cells distinct tonoplast localization transiently appears before cells enter the elongation zone (Fig. 5 C,E, Supplementary data Video S5 of FH1-GFP-labelled tonoplast in transition zone cells with small vacuoles, Supplementary data Video S6 of FH1-GFP-labelled tonoplast in older transition zone cells with a large central vacuole; both videos are from *fh1-4 pFH1:FH1-GFP* plants, acquired under the same conditions as the images shown in Fig. 5 A, scale bar = 10 µm.). Further up the root, in the elongation zone, FH1-GFP-positive mobile particles disappear and signal intensity gradually diminishes until no signal can be detected as the maturation continues (Fig. 5 C).

Plasmalemma localization of FH1-GFP in meristematic cells was confirmed using FM4-64 co-localization that also documented that the formin is targeted already to the nascent cell plate (Fig. 5 F).

### FH1 targets a subset of endomembranes whose organization is actin-dependent

To examine the nature of the intracellular FH1-GFP positive compartments, we first used the FM 4-64 membrane staining (Fig. 6 A). In the apical meristem, some of the mobile formin-positive particles took up the dye as early as after 15 minutes and represented thus early endosomes, but the majority acquired FM4-64 signal later, consistent with FH1-GFP labelling late endosomes. The kinetics of FM4-64 labelling of the formin-positive compartments indeed suggests that FH1 associates with at least two types of endomembrane compartments – a fast-staining subset of early endosomes and a larger population of late endosomes (Fig. 6 B, Supplementary data video S7 documenting a subset of mobile FH1-GFP-positive particles stained with FM4-64 after 80 min of incubation with the dye, acquired under the same conditions as the images shown in Fig. 6 A; scale = 10 µm.).

**Figure 6.**
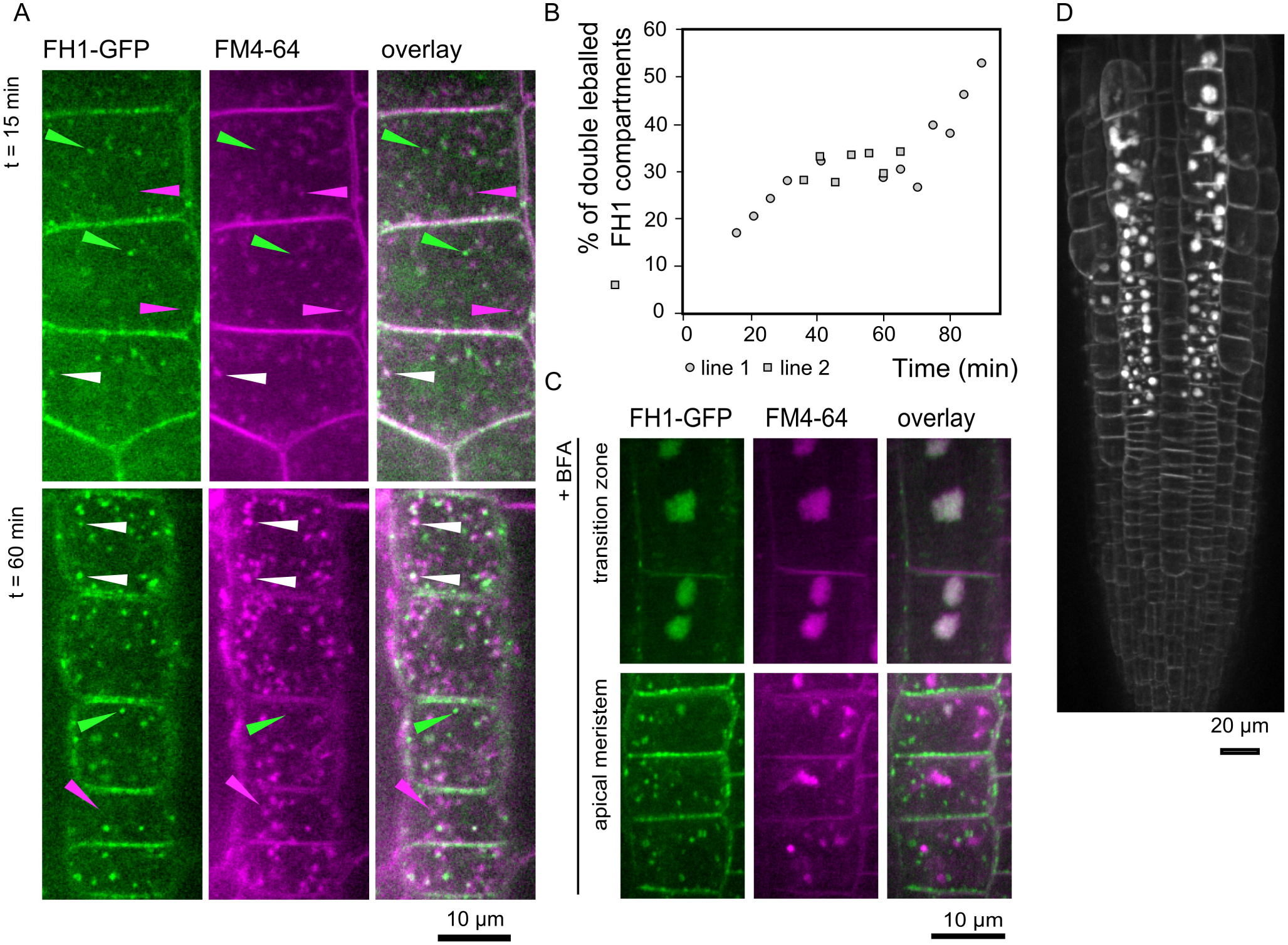
In roots, FH1-GFP associates with late endosomes and BFA-sensitive compartments. (A) Root apical meristem of *fh1-4 pFH1:FH1-GFP* plants stained with FM4-64 for 15 min to label early endosomes and for 60 min to visualize late endosomes. White arrowheads: FH1-GFP positive endosomes, green and magenta arrowheads: structures with GFP or FN4-64 signal only, respectively. (B) Time course of FM4-64 uptake into FH1-GFP positive compartments in the root apical meristem of seedlings derived from two transgenic lines from the progeny of the cross between *fh1-4 pFH1:FH1-GFP* and WT. Percentages of compartments positive for GFP, FM4-64 and both markers at given time points are shown. Data points indicate average from at least 100 dots from a minimum of 5 cells. The fraction of doubly labelled particles is probably underestimated due to a delay in green and red chanel capturing, combined with particle mobility. (C) Apical meristem and transition zone after incubation with 40 μM BFA. Seedlings were pre-stained with FM4-64 to visualize endosomes. Z-projections of a confocal laser scanning microscopy (CLSM) stack are shown. (D) Endosome aggregation in the root tip of an *FH1-GFPox* plant (Z-projection of a CLSM stack). Images from 5 days old seedlings.

After application of Brefeldin A (BFA), an inhibitor of vesicle transport, the plasmalemma signal and static particles remained unchanged, but the mobile FH1-GFP particles in the transition zone aggregated to typical BFA bodies. However, in the apical meristem, where only a part of endomembrane compartments collapsed into BFA bodies under identical treatment conditions, FH1-GFP-positive structures did not aggregate, suggesting that the formin is associated with a subset of endomembranes less sensitive to BFA (Fig. 6 C). Interestingly, structures reminiscent of BFA bodies were also observed, alongside numerous small mobile endosomes, in the root tips of plants from the incidental overexpression line *FH1-GFPox* even without BFA treatment (Fig. 6 D).

We further tested if the organization of the FH1-GFP-positive compartments depends on the actin cytoskeleton. Treatment with Latrunculin B (LatB) in a manner substantially disrupting the actin network (Supplementary data Figure S8) had no effect on the plasmalemma-associated signal but resulted in marked enlargement, or aggregation, of the FH1-GFP positive compartments in both root transition zone (Fig. 7 A) and apical meristem (Fig. 7 B). At the same time, FH1-GFP particles came to a halt. In the transition zone, vacuolar fragmentation was observed, as reported previously (Scheuring et al. 2016). Cells that were at the transient tonoplast labelling stage during LatB treatment exhibited variable intensity of the FH1-GFP signal in distinct sub-vacuoles (Fig. 7 C).

**Figure 7.**
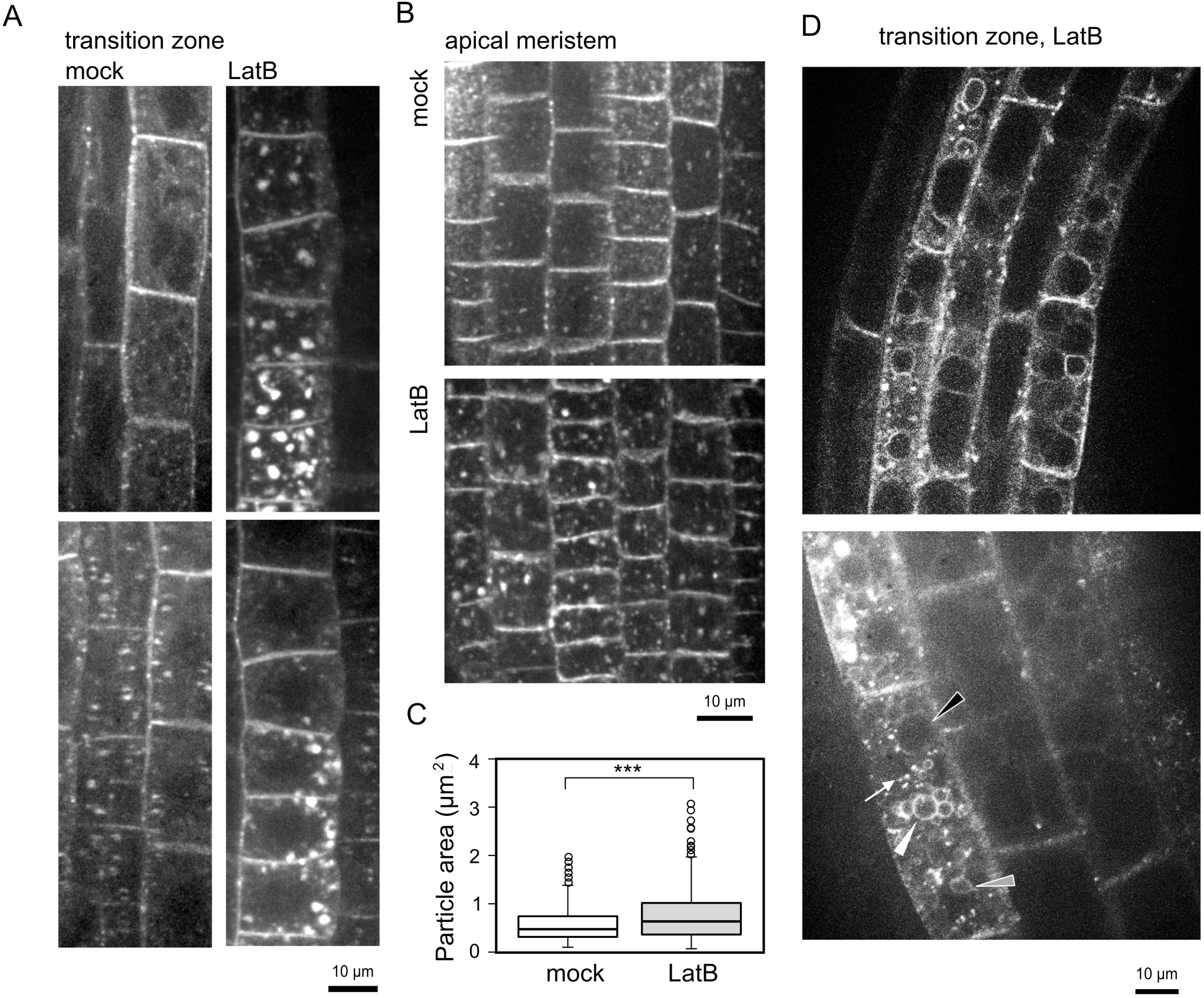
Localization of FH1-GFP in roots depends on microfilaments. (A) Effect of 3 hours of treatment with 10 μM LatB on different root tip zones of *rdr6-12 pFH1: FH1-GFP* seedlings, compared to mock treatment (DMSO). Z-projections of two views on the transition zone, periclinal membranes to the right. (B) The same treatment and genotype as in (A), Z-projection of a view of the apical meristem. (C) Size distribution of FH1-GFP positive compartments in *fh1-4 pFH1:FH1-GFP* plants treated as in (A) and (B). For each treatment at least 170 particles were measured. Triple asterisks denote a highly significant difference (t-test P < 0.001). (D) Vacuolar fragmentation in the transition zone of *fh1-4 pFH1:FH1-GFP* roots treated as in (A) to (C). Top: a single CLSM optical section. Bottom: Z-projection. Note the presence of vacuoles with strong, weak and no FH1-GFP signal (filled arrowheads in shades from white to black), as well as non-vacuolar FH1-GFP-positive bodies (arrow), within the same cell All images are from 5-6 days old seedlings.

We can thus conclude that the organization and dynamics of the FH1-decorated endomembrane compartments requires an intact actin cytoskeleton.

## Discussion

Typical plant Class I formins, including FH1 (At3g25500), the most ubiquitously expressed member of this formin clade in Arabidopsis vegetative tissues (Hruz et al. 2008), are predicted to be integral membrane proteins (Cvrčková 2000). The presence of extensin-like motifs in the N-terminal extracytoplasmic domain suggests cell wall attachment, and indeed FH1 and its relatives are generally believed to be targeted to the plasmalemma (e.g. van Gisbergen and Bezanilla 2013). Such localization is supported by observations from heterologous overexpression of FH1 in mature tobacco leaf epidermis (Martiniere et al. 2011) and in tobacco pollen tubes (Cheung and Wu 2004), and documented also for several related Class I formins (see Introduction).

We employed transgenic Arabidopsis expressing biologically active C-terminally GFP-tagged FH1 (FH1-GFP) from its native promoter to investigate *in vivo* localization of this protein. Trichome shape alterations in plants incidentally overexpressing FH1-GFP indicated possible participation of FH1 in trichome development. Alternative explanations, especially transgene-induced mutagenesis, while not excluded, are less likely, since the trichome deformity appears to be dominant. We were so far unable to successfully overexpress FH1-GFP in stably transformed Arabidopsis under a constitutive promoter, nor were stable overexpression lines reported by others; thus, the final proof of causal connection between FH1 overexpression and trichome deformation is still lacking .Nevertheless, consistent with FH1 contributing to trichome morphogenesis, WT plants, *fh1* mutants and plants overexpressing FH1-GFP exhibited an inverse correlation between the level of FH1 expression and number of trichome branches, as well as the extent of epidermal pavement cell lobing. It is thus tempting to speculate that FH1 may generally act to control cell shape complexity, as suggested also by previous observations in pavement cells (Rosero et al. 2016).

Surprisingly, although FH1-GFP resides mainly at the plasmalemma in fully differentiated epidermal cells and is present at the plasmalemma also in root tissues up to the elongation zone, the protein relocates between several membrane compartments in expanding pavement cells and in immature root tissues. In such cells, FH1-GFP is found also in mobile membrane compartments within the cytoplasm, and in the root transition zone it transiently decorates the tonoplast at or around the stage when small vacuoles fuse to generate the large central vacuole (see e.g. Kolb et al. 2015). The protein localization pattern was generally independent of the allelic status of *FH1* and *RDR6* (Supplementary data Table S9, Supplementary data Figure S10), a gene whose mutation prevents transgene silencing, but has also additional phenotypic effects (Peragine et al. 2004). Our findings raise several questions. What is the identity of the FH1-positive compartments, how is the formin targeted into them in a cell type-specific manner, and what is the biological role of the complex and dynamic relocation of FH1 during cell differentiation?

While plasmalemma-localized proteins pass through the secretory pathway, it is unusual for a transmembrane protein to dwell in several distinct membrane compartments depending on the differentiation status of the cell. Very few dual- or multiple-localized plant membrane proteins have been reported, most of them targeted to secretory pathway compartments, plastids and/or mitochondria (see Kasaras and Kurze 2017). Some SNAREs can localize to either the endoplasmic reticulum or the plasmalemma (Suwastika et al. 2008). However, none of the observed FH1-positive intracellular compartments resembles the well known architecture of mitochondria, plastids or the cortical endoplasmic reticulum. Dual localization between the tonoplast and the trans-Golgi network has been reported for Arabidopsis AtPAT10 S-acyl transferase, but, unlike FH1, this protein was not found on the plasmalemma (Qi et al. 2013). Dual tonoplast and plasmalemma localization of DMP1, a membrane protein of unknown function, results from the use of alternative translation initiation sites generating two protein isoforms capable of inserting into the membrane (Kasaras and Kurze 2017). However, in FH1 an alternative translation start at the 2nd in-frame ATG would eliminate the signal peptide. Remarkably, in various plant species a fraction of some tonoplast proteins can be detected at the plasmalemma (e.g. Robinson et al. 1996, Nilsson et al. 2010). Some Arabidopsis tonoplast aquaporins (TIPs) transiently label the plasmalemma during seed maturation before settling at their tonoplast destination in mature seeds (Gattolin et al. 2011), thus exhibiting a pattern inverse to that seen for FH1 whose transient tonoplast localization in the root transition zone precedes plasmalemma localization in mature tissues.

Membrane proteins co-translationally inserted into the endoplasmic reticulum can reach the tonoplast by multiple routes. Vacuolar targeting depends on the presence of specific motifs within the cytoplasmic portion of the protein (Pedrazzini et al. 2013). Indeed, the cytoplasmic part of FH1 contains multiple L-L and one F-Y motif that might confer tonoplast localization, albeit all but one of the di-leucine motifs are part of the conserved FH2 domain and likely to be obscured by its folding. Vacuolar targeting may be controlled by cell type-specific protein-protein interactions or by post-translational modifications; such mechanisms may also account for the other observed localization changes. Indeed, FH1 can undergo phosphorylation (Curran et al. 2011).

Cell type-specific localization might also result from alternative mRNA splicing generating multiple protein isoforms. However, the current *A. thaliana* genome annotation (Krishnakumar et al. 2017) only predicts a single FH1 mRNA comprising four exons. Also all empirically obtained cDNA sequences derived from the At3g25500 locus found in the GenBank database match the predicted splicing pattern, or at least are consistent with it (Supplementary data Figure S11). While two alternative transcripts are recorded in the Arabidopsis Reference Transcript Dataset (ARTD2, Zhang et al. 2017), these only differ in the position of the transcription start site and encode identical proteins. We thus consider alternative splicing unlikely, although we cannot entirely exclude this possibility.

In immature epidermal pavement cells, in root apical meristem and in the root transition zone cortex, FH1-GFP decorated cytoplasmic particles that exhibited actin-dependent mobility. As confirmed by staining with the membrane dye FM4-64, these particles are endomembrane compartments or aggregates thereof. The kinetics of dye uptake is comparable to that of FM1-43 in late endosomes (Emans et al. 2002). Surprisingly, the uptake curve is biphasic, suggesting two endosomal populations positive for FH1-GFP, one taking up the dye readily, and the other after a delay, which, however, might also merely reflect changes in the cell physiology due to submersion in the staining solution. In any case, some of the mobile particles in the root apical meristem do not readily incorporate freshly endocytosed membrane, and may be of a late-endosomal or non-endosomal character. Interestingly, the pathogen-responsive Class I formin FH4 was observed not only in distinct plasmalemma domains adjacent to the invading pathogen (including immobile dots at sites of callose deposition), but also in mobile intra-cytoplasmic particles whose majority was distinct from early endosomes (Sassman et al. 2018).

Further clues to the identity of the FH1-positive compartments may be obtained from application of the exocytotic membrane trafficking inhibitor Brefeldin A (BFA) that causes reversible collapse of the endoplasmic reticulum and the Golgi apparatus into a “BFA compartment” (Ritzenthaler et al. 2002). In the root transition zone most mobile cytoplasmic FH1-GFP particles fuse with the BFA compartment and the tonoplast signal disappears, indicating that FH1 is transported to the tonoplast via the BFA-sensitive import pathway (Pedrazzini et al. 2013). However, in the apical meristem only a small fraction of BFA compartments carries FH1-GFP, while most FH1-GFP-labelled compartments are BFA-resistant, as expected for endosomes. Thus, FH1, depending on the cell differentiation status, localizes to (at least) two types of endomembrane compartments, reminiscent e.g. of the Arabidopsis flagellin receptor that can be also found in either BFA-sensitive or BFA-resistant compartments depending on its activation status (Beck et al. 2012). An alternative explanation involving differences in sensitivity of various root tip cell types towards BFA, as reported e.g. for maize (Baluška et al. 2002), cannot be excluded. Nevertheless, the near-absence of FH1-GFP from BFA compartments that do form in the root apical meristem (albeit less easily than in the transition zone) indicates that FH1 decorates a specific subset of endomembranes whose behaviour, if not identity, changes during cell differentiation.

What function could endomembrane-localized FH1 perform? We previously showed that *fh1* mutants exhibit decreased mobility of clathrin light chain (CLC)-positive compartments (Rosero et al. 2016). While these compartments may represent endosome aggregates rather than individual endosomes, localization of the formin to endosomes may provide a mechanistic basis for the observed altered mobility. However, for a definite proof of FH1 participating in endosome transport, a detailed quantitative study of endosome kinetics in WT and mutant plants would be required. FH1 may also participate in development of the large central vacuole of differentiated root cells, since biogenesis of the central vacuole during root development is both actin- and microtubule-dependent (Zheng et al. 2014), and actin also mediates auxin-controlled vacuole expansion in late root meristem/transition zone (Scheuring et al. 2016)., Alternative explanation that FH1 incorporates into the tonoplast during membrane recycling is possible, although in such a case transient tonoplast labelling would be expected to occur in various cell types during development, which was not observed.

Incidental overexpression of FH1-GFP induces formation of large endomembrane bodies in transition zone cells, reminiscent of BFA bodies or the enlarged endomembrane compartments found in plants defective in the exocyst complex mediating targeted exocytosis (Drdová et al. 2013). Enlargement of FH1-GFP-positive compartments was also observed upon disruption of microfilaments, indicating an intimate link between actin cytoskeleton and endomembrane architecture. Indeed, actin disruption upon BFA treatment is known in plants (Alvarez and Sztul 1999, Hörmanseder et al. 2005, Zhang et al. 2010), and changes in endomembrane system organization upon perturbation of microfilaments have been reported in pollen tubes (Zhang et al. 2010, Moscatelli et al. 2012). Class I formins such as FH1 are prime candidates for linking endomembranes to microfilaments, and the overexpression phenotype appears to support this hypothesis.

In lobed epidermal pavement cells, i.e. at a more advanced stage of differentiation, FH1-GFP locates predominantly to the plasmalemma, and is excluded from microtubule-occupied areas, as reported previously (Martiniere et al. 2011). However, distribution of the formin at the plasmalemma is heterogenous, with distinct foci in actin-rich cortical regions. Our observations were made in 6 days old plants, whose cotyledon epidermal pavement cells are still growing (Rosero et al. 2016), but already past the stage of cell lobe initiation (Armour set al.2015). The localization of the formin at that stage is thus obviously irrelevant to cell shape determination. In younger pavement cells we observed some heterogeneity in the FH1-GFP fluorescence intensity, but we were so far unable to determine whether it predicts the future pattern of cell lobing. In developing trichomes of true leaves, however, FH1-GFP decorates the plasmalemma at expanding cell tips.

The FH1-GFP-labelled plasmalemma foci co-localize with a plasmodesmal marker in the cotyledon and with callose deposits in developing root tissues, consistent with a very recent report (Diao et al. 2018), published while this study was being prepared for submission, of similar targeting of a C-terminally truncated FH1 (residues 1-587), found at plasmodesmata of aboveground organs when overexpressed from the 35S promoter. Remarkably, Diao et al. (2018) do not discuss their observations in the context of a previous study (Martiniere et al. 2011) that documented diffuse plasmalemma localization with exclusion of microtubule-occupied regions for full-length FH1, and uniform plasmalemma decoration for its shorter deletion derivative (residues 1-140), in transient heterologous expression experiments. However, while integral components of the plasmodesmata stay in the plasmalemma after plasmolysis (Thomas et al. 2008), some of the FH1-GFP signal loses plasmodesmal localisation in plasmolysed cells except in the plasmalemma regions that remain attached to the cell wall, resulting in weakening of the foci. Thus, while FH1-GFP remains at the plasmalemma under these conditions, its association with plasmodesmata may be, at least for a subpopulation of the protein, rather loose and possibly related to microfilaments, which are enriched in plasmalemma domains resisting plasmolysis (Lang et al. 2014). Our findings suggest that alterations in FH1 activity may influence the function of plasmodesmata. Indeed, loss of function *fh1* and *fh2* mutations increase plasmodesmal permeability in leaves, with an additive effect in double mutants (Diao et al. 2018). Since formins participate in the organization of cell to cell junctions also in metazoan systems (Grikscheit and Grosse 2016), it is tempting to speculate about their possible ancient role in intercellular communication.

The complex, dynamic relocation pattern of FH1 has not been observed until now, possibly for methodological reasons. Knowledge on FH1 localization comes from cell fractionation that only captured the difference between membrane and soluble fractions (Banno and Chua 2000), from transient heterologous expression in mature tobacco pavement cells (Martiniere et al. 2011) and in pollen, where little if any of the protein is naturally expressed (Cheung and Wu 2004), driven by a strong promoter (bringing added risk of overexpression artefacts). Use of the natural promoter and stable expression lines enabled us to follow the fate of biologically active protein throughout the development, revealing a surprising richness of localization. Our observations may reflect the presence of cell type-specific interaction partners and/or post-translational modifications, possibly resulting in the existence of distinct developmental windows enabling protein incorporation into structures such as the plasmodesmata, inaccessible to study using transient expression assays. Besides of bringing new knowledge on the functioning of the Class I formin FH1, our present study thus also underlines the benefits of stable expression from natural promoters in plant protein localization studies.

## Materials and methods

### Plants

Previously described *A.thaliana* lines carrying Lifeact-mRFP, mCherry-TUA5 (Rosero et al. 2016, Cvrčková and Oulehlová 2017) and the At5g24010-RFP (Fernandez-Calvino et al. 2011), as well as *FH1* WT and *fh1-1* mutants (Rosero et al. 2013) were employed. To detect WT *FH1* and *fh1-1* alleles, PCR was used as described previously (Rosero et al. 2013). The *fh1-4* mutant line corresponds to the SALK T-DNA insertional line N051065. Presence of the *fh1-4* insertion was verified by PCR using primers SALK LBb1.3 (5’ ATTTTGCCGATTTCGGAAC 3’) and fh1-4RP (5’ CCAAGCTTAACAGGCGAAT 3’), while for the WT allele primers fh1-4LP (5’ GAGTCAGGTGACTACTAAAGCG 3’) and fh1-4RP were used. Absence of full-length *FH1* transcript in *fh1-4* mutant was confirmed by RT-PCR as described previously (Rosero et al. 2013) using primers 5FH1for (5’ CGAAGAAGCAAACGTAAC 3’) and 5FH1rev (5’ AGAGCCTCACAAACTTCTTCTAT 3’) for detection of 5ill terminal region, as well as 3FH1for (5’ CAGTAGGATGTTCTTAAAGCT 3’) and 3FH1rev (5’’ AGACAAAGCTGAGAGCGGCCGCGAAGAA 3’) for detection of 3ill terminal region of the FH1 transcript. As a control, *ACT7* transcript was amplified using primers ACT7F (5’ GCCGATGGTGAGGATATTCAGC 3’) and ACT7R (5’ CAAACTCACCACCACGAACCAG 3’). Decrease of pavement cell circularity comparable to that observed in *fh1-1* was also documented for *fh1-4* (Supplementary data Figure S12). For some experiments, *rdr6-12* plants (NASC line CS24286, Sasse et al. 2015) were used. All plants were in the Col-0 background.

For propagation, crosses, transformation and trichome morphology analyses, plants were grown in peat pellets (Jiffy) under long day conditions. Seedlings for imaging were grown *in vitro* at 22 °C with a 16 h light/8 h dark cycle on vertical half strength Murashige and Skoog (MS) plates from surface-sterilized seeds stratified by several days of post-imbibition storage at 4°C prior to sowing; seedling age is counted from plating.

### Cloning and plant transformation

The At3g25500 genomic locus including introns and 2.8 kbp upstream from the start codon was amplified using Phusion polymerase (New England BioLabs, Ipswich, MA, USA) and primers FH1-fw (5’ CGCGGATCCCATTCATATGAAAAGAGTTGACC 3’) and FH1-rv (5’ TATGCGGCCGCAAAGAAACTAATGAGATTGAGTTATGTTC 3’), cloned into the Gateway pENTR3C vector and verified by sequencing. The entry clone was transferred into pGWB4 destination vector (provided by Tsuyoshi Nakagawa, Shimane University, Japan) using LR clonase (Invitrogen) and the resulting FH1:FH1-GFP construct was introduced into *Agrobacterium tumefaciens* and transformed into *A. thaliana* using the floral dip method. Transformants were selected using hygromycin and kanamycin.

### Staining and drug treatments

For plasmalemma staining of aboveground organs, seedlings were incubated for 15 min with 5 μM FM4-64 (Invitrogen, storage stock in water) in water. For the first 5 min, vacuum was applied to enhance dye penetration. For the plasmolysis experiment, seedlings after plasmalemma staining were rinsed in water and further incubated with 1M mannitol water solution. For root staining, liquid ½ MS with 5 μM FM4-64 was applied without vacuum treatment for 10 min to stain the plasmalemma and for 15 - 90 min for endosome visualization. Seedlings were rinsed in water before microscopy.

For callose detection, 4 days old seedlings were stained with aniline blue solution (0.5 mg/ml anilin blue dissolved in deionized water) for at least 1 hour in dark conditions. After staining, seedlings were rinsed in water to remove excess aniline blue and examined by confocal microscopy.

For BFA treatment, seedlings were first stained for 15 min with FM4-64 as described above, rinsed in water and subsequently incubated in dark with 40 μM Brefeldin A (Sigma, storage stock in dimethylsulfoxide – DMSO) in water. For actin depolymerization assay, seedlings were transferred from plates to liquid ½ MS media with Latrunculin B (LatB) of final concentration 10 μM (Sigma, storage solution in DMSO). 1% DMSO in liquid ½ MS was used for control (mock) treatments.

### Microscopy

For routine evaluation of transgenic plant fluorescence and for capturing bright field images, an Olympus BX-51 microscope equipped with an Olympus DP50 camera has been used. Olympus Provis AX 70 with a 20x water-immersion objective was used for visual trichome branch counting, which was performed on detached first true leaves from 24 DAG plants that were prior to observation cleared for 6 days in a solution prepared by mixing 120 g chloral hydrate, 7.5 ml glycerol and 150 ml of water. All trichomes found on one half of at least 5 leaves per genotype were evaluated.

Confocal laser scanning microscopy (CLSM) and spinning disk confocal microscopy (SDCM) images were acquired as described previously (Cvrčková and Oulehlová 2017). For CLSM, a Zeiss LSM880 microscope with a 63x/1.2 water-immersion objective and 488-nm argon laser for excitation, with optical slice thickness set at 0.7 to 1 µm, was used. SDCM recordings were performed using a Yokogawa CSU-X1 inverted spinning disk confocal microscope on a Nikon Ti-E platform, with laser box Agilent MLC400, camera Andor Ixon, objective plan apochromat × 100 oil (NA = 1.45), laser lines set at 488 and 561 nm, and image interval 1 s.

### Image analysis

Pavement cell size and circularity was determined using manual tracing as described previously (Rosero et al. 2016). In some experiments, a modified protocol, which produced results fully co-measurable with manual tracing, was used as follows. One stack of three CLSM images of adaxial epidermis from the apical third of a cotyledon stained with 1μM FM4-64 was taken per plant from total of 10 plants per genotype, and Z-projection was generated for each stack using Fiji (Rueden et al. 2017). The image was then processed in Fiji applying first smoothing and then thresholding using the Huang algorithm manually adjusted to achieve good cell contour detection. The resulting image was converted to binary and skeletonized, gaps in cell contours were manually closed, and cell area and circularity measured using Fiji tools for total of 8–10 cells per sample within a field. To avoid bias, cells that made contact with a diagonal line were chosen in each field.

To quantify FM4-64 internalization into FH1-GFP positive compartments, we collected single CLSM optical sections from plants stained with FM4-64 for each timepoint. For at least five cells per sample, the total number of GFP-positive particles in the plane of focus was determined, together with the fraction of particles that were also FM4-64 positive.

FH1-GFP compartment size was estimated by measuring the cross-section area of fluorescent particles in Fiji in single confocal sections after manual particle perimeter tracing. For each treatment, three optical sections, each taken from a different plant, were evaluated by measuring all particles found in 10 cells per section.

### Funding

This work was supported by the Grant Agency of the Czech Republic [grant 15-02610S] and Ministry of Education, Youth and Sports of the Czech Republic [project NPUI LO1417]. The microscopy facilities used in this work were supported by the European Regional Development Fund and the state budget of the Czech Republic [projects CZ.1.05/4.1.00/16.0347 and CZ.2.16/3.1.00/21515], and by Ministry of Education, Youth and Sports of the Czech Republic [project LM2015062 Czech-BioImaging].

### Disclosures

No conflicts of interest declared.

## Acknowledgements

We thank Jana Šťovíčková and Marta čadyová for technical assistance.

